# Embryo spatial 3D genomics

**DOI:** 10.1101/2024.05.07.592900

**Authors:** Yuang Ma, Bo Gou, Yuetong Xu, Muya Shu, Falong Lu, Xiang Li

**Affiliations:** State Key Laboratory of Plant Genomics, Institute of Genetics and Developmental Biology (IGDB), Chinese Academy of Sciences, Beijing 100101, China; University of Chinese Academy of Sciences, Beijing 100049, China

**Author notes:** Corresponding (X.L.). These authors contributed equally to this work (Yuang Ma, Bo Gou, Yuetong Xu).

## Abstract

The 3D architecture of the genome is crucial for controlling gene expression and organ development. Here, we introduce a spatial 3D genomics approach for assessing chromatin conformation *in-situ* in tissue sections, by integrating microfluidic deterministic barcoding and SPRITE procedures. This method was applied to mouse embryo sections, revealing a hierarchical model of chromatin interactions within and between compartments in various organs. The intra-compartment interactions vary among organs to orchestrate gene expressions, while the inter-compartment interactions remain identical in the most organs. Beyond this, the liver exhibits overwhelmingly packed chromatin with enhanced adjacent-compartment interactions, possibly related to its physiology. These findings highlight the importance of tissue-spatial information in understanding embryonic chromatin organization. The approach presents a powerful tool for investigating these processes in tissues with high heterogeneity.

**One Sentence Summary:** A spatial 3D genomics approach was developed that accesses hierarchical chromatin conformation *in-situ* in tissue sections.

---

In eukaryotes, DNA undergoes regular folding and precise organization with chromatin proteins, forming functional hierarchical conformations (*1-4*). Advances in single-cell 3D genomics approach have enabled the assessment of how these conformations affect cell cycling and aging trajectories (*5*). However, when applying to isolated cells in interested organs, these approaches struggle to fully access the tissue development or lesion, as they cannot obtain the tissue-spatial information of detected cells. We introduce a pioneering spatial 3D genomics technology that comprehensively assesses chromatin conformation for each tissue spot on an entire section. The method integrates two procedures: tissue-spatially barcoding and genome architecture capturing. Two classes of spatially barcoding systems were previously developed: one utilizes adaptor-ligated arrays, as seen in Slide-seq (*6*), Stereo-seq (*7*), and Visium platforms, while the other uses parallel microfluidic channels delivering unfixed adaptors into the nucleus on tissue sections (*8*), as in Spatial-CUT&TAG and ATAC for collecting DNA information in the nucleus (*9, 10*). For genome architecture capturing, techniques such as Hi-C (*8*) and Dip-C (*9, 10*) isolated one cell into each PCR well and sequenced proximally-ligated DNA ends. In contrast, the SPRITE approach performs split-pool barcoding of DNA end complexes to identify multi-way interactions of thousands of individual cells (*11*). In this study, we design a Spatial-SPRITE approach, combining SPRITE barcodes with DNA end complexes ligated by microfluidic-based spatial barcodes (Fig 1A). Using the Spatool algorithm pipeline, an embryonic atlas of chromatin conformation at a high tissue-spatial resolution was constructed, offering unprecedented insights into interactions within and between compartments in organ regions and sub-regions.

**Fig. 1.**
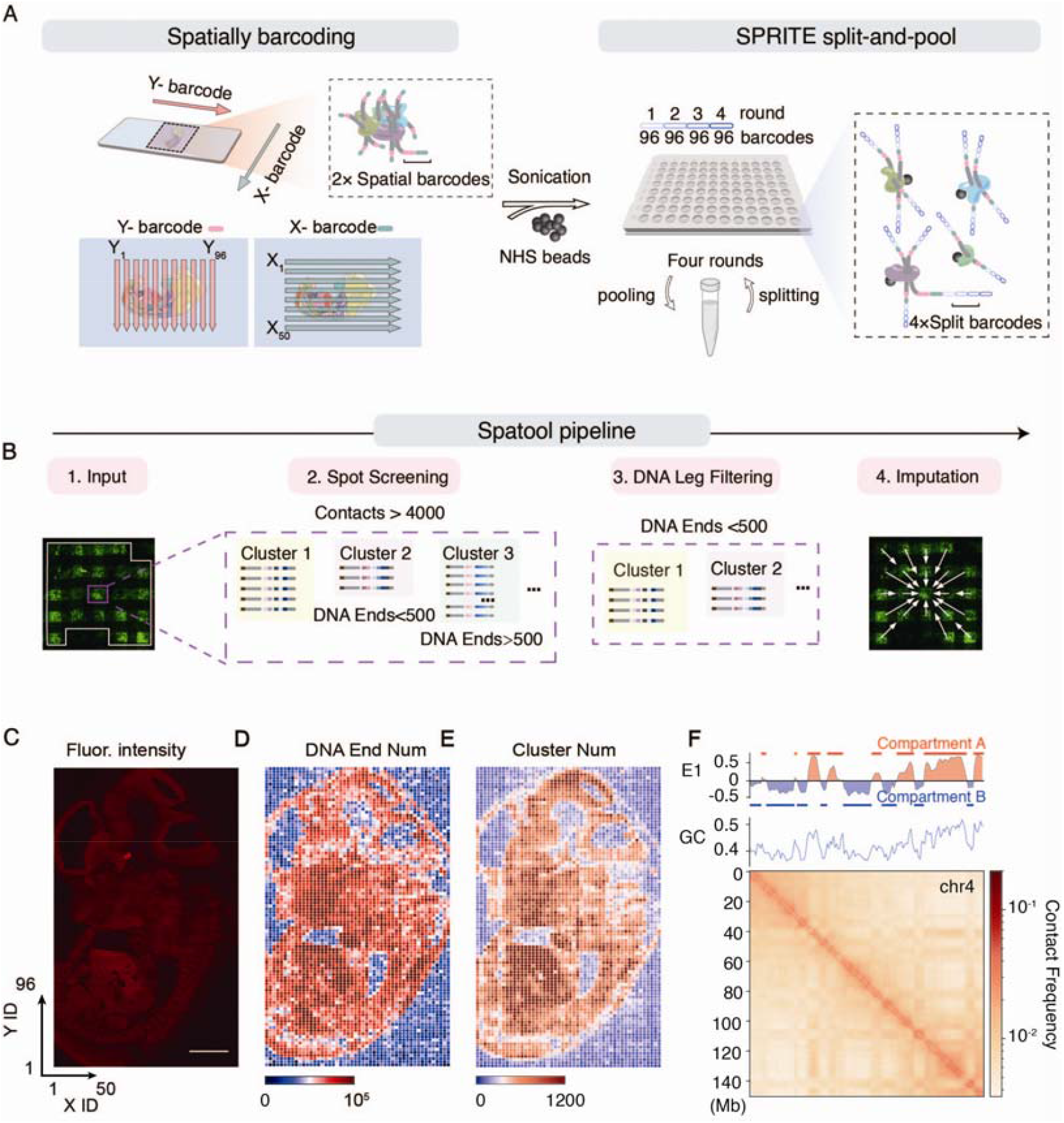
Diagram of spatial-SPRITE experimental procedure and Spatool pipeline. (**A**) Schematic of spatial-sprite workflow. (**B**) Verification of inter-channel closure. The E10.5 mouse embryo section was added with TAMRA-labeled X-direction barcode (left) and FAM-labeled Y-direction barcode (right). Magnified image showed that there is no leakage among channels during 1streaction or 2nd reaction. Scale bars, 500 µm or 250 µm (zoomed). (**C**) Fluorescence results of mouse embryo. (**D**) The heatmap shows the number of DNA ends within each Spot. (**E**) The heatmap shows the number of clusters within each Spot. (**F**) Spatial-SPRITE derived chromatin contact maps on chromosome 4 at 1◻Mb resolution.

## In the spatial bSpatial-SPRITE procedure design

In the spatial barcoding process (Fig S1A), tissue section was firstly prepared on a slide. Within the nucleus, DNA and proteins were crosslinked, and then DNA was cleaved to expose the ends for subsequent barcoding. We found that using double restricted enzymes (MseI & HinP1I) is more efficient than DNase I for cleavage, as shown by distinct fluorescent signals after fluorescence-labelled adaptor ligation (Fig S1B). Next, we designed polydimethylsiloxane (PDMS) microfluidic chips with 96 horizontal and 50 vertical channels, each 50 μm wide (Fig S2A-C). The tissue section was covered with a horizontal-channel chip, allowing Y_1_-Y_96_ barcodes to flow and ligate to tissue DNA ends in the channels. Cleaning the section, X_1_-X_50_ barcodes were also ligated to these ends using a vertical-channel chip. The resulting fluorescent bands and spots on sections reveals Y_n_ and X_n_ barcode-labelled tissue (Fig S2B).

In the SPRITE process (Fig S1A), the tissue section was firstly fragmented into DNA-protein complexes using ultrasonication. To ensure valid interaction information, we tested different sonication durations and found that ten minutes provide the more complexes with 10-1000 DNA legs (Fig S3A-B). Following ultrasonication, the complexes are coupled with NHS-activated beads, and rounds of split-pool processing were used to ligate each single DNA-protein complex to a unique barcode combination. This is achieved by splitting the beads suspension into 96 PCR wells for each round, where distinct DNA barcode tags are contained. We performed four rounds of split-pooling and counted the complex identities relying on one to four split barcodes of sequencing reads (Fig S3C). The results show that decreasing rounds may lead to insufficient barcode combinations labeling complexes. Subsequently, the DNA ends combined with spatial and SPRITE barcodes, undergo library preparation, including ligating one-side sequencing adaptors and adding the other side through random primer annealing and extending to ensure an appropriate sequence length (350 to 750bp).

## Spatool pipeline processes Spatial-SPRITE sequencing data

We conducted Spatial-SPRITE on two sections of E12.5 mouse embryos, yielding 3.5 billion sequencing reads (Table S1). The reads contained all six spatial and SPRITE barcodes (Fig S4A), and over 50% (746 million and 1026 million) of reads showed complete barcode sets (Fig S4B), indicating high data quality. The non-duplicated reads (DNA ends) reached a saturation point (Fig S4C), suggesting the additional sequencing may not significantly increase the depth or quality of the data. To analyze the data, we developed the Spatool pipeline (Fig 1B), which included tissue spot screening, DNA complex filtering, and data imputation. In the all 4800 (96 Y-× 50 X-) pixels, we identified 3133 tissue spot pixels with over 4000 DNA legs for analysis, with some pixels containing up to 100,000 DNA ends. These pixels lie in a section-like shape (Fig 1C-E). We also detected approximately 95% of DNA ends in these valid pixels (Fig. S5A), confirming the reliability of the Spatial-SPRITE experiments. Complexes with over two DNA ends were considered valid for estimating DNA contacts, representing about 30% (Fig S4D), a proportion consistent with previous reports (*12*). We then removed the data of complexes containing over 500 DNA ends, as they may not have been fragmented sufficiently when ultrasonication. Using E1 values, we identified compartments A and B of whole embryo chromatins, known as loose and condensed regions, respectively, with high and low gene expression(*13*). Heat maps of chromatin interaction confirmed these boundary-clear compartments (Fig 1F). The E1 value curve correlated with the GC contents of bins (Fig 1F), consistent with previous report (*14*). To further distinguish the chromatin conformation of each organ, we enhanced the distinction of tissue spot pixels by imputing DNA ends from the adjacent pixels.

## A high-resolution chromatin conformation atlas for embryonic organs

Following the Spatool processing, the pixels for section spots (Table S2) were clustered using 200 principal components of DNA ends per bin, explaining over 80% of the interpretability variance (Fig S6A). This resulted in 30 pixel clusters. Assigning pixels with their spatial coordinates (X- and Y-ID) revealed a chromatin conformation atlas of the 13 previously defined organs (*7*) (Fig 2AB; Fig S6BC). Interestingly, some organs like the brain and cartilage primordium were split into sub-regions, suggesting that the divergence of developmental fates for these organs precedes this E12.5 stage. Previous reports applying single cell 3D genomics approaches, isolated the defined cerebellum or cerebral cortex of human and mouse (*15*) to assess varying chromatin conformations of brain sub-regions. However, our study detected nine sub-regions in the brain, beyond the concepts of embryonic fore-, mid-, and hind-brains, providing a new insight into the heterogeneity during brain differentiation and development.

**Fig. 2.**
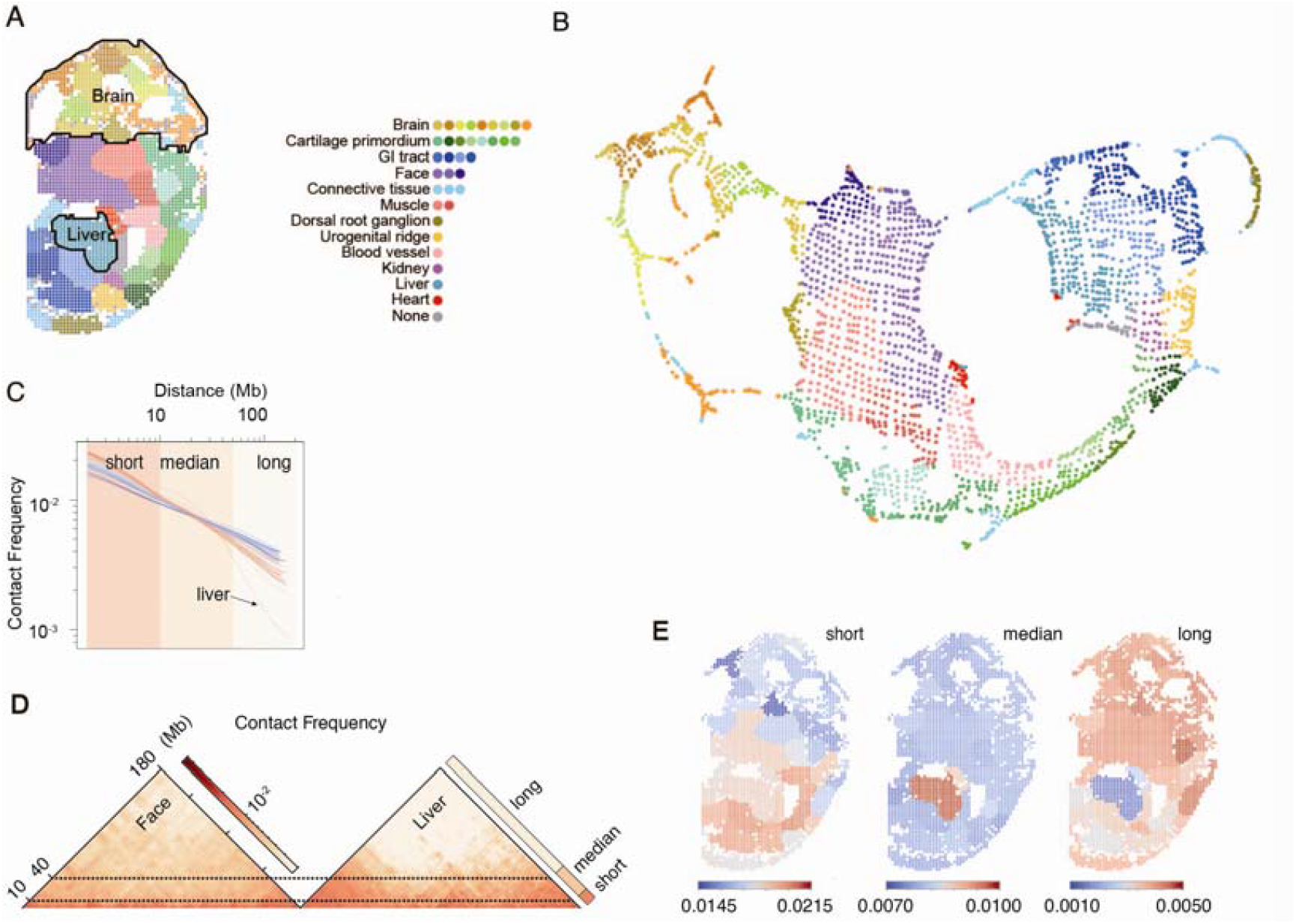
UMAP plot and spatial distributions of tissue spot pixels. (**A**) The result of k-means algorithm uses the interaction information to cluster the spot. (**B**) The interaction information is mapped to 2D space via UMAP. The different colors represent different classes of spatially corresponding tissues. (**C**) The relationship between the number of interactions and the distance of interactions. (**D**) Comparison of chromosome 4 contact maps of face and liver. (**E**) The frequency of interactions between tissues at different distances.

## Embryonic organs exhibit hierarchical chromatin conformations

Furthermore, based on this atlas, we detected hierarchical conformations for short-(less than 10 Mb), median-(10 to 40 Mb apart), long-distance (over 40 Mb) DNA interactions, which unexpectedly vary among organ regions and sub-regions (Fig 2C-E). These hierarchical conformations may not be identical to the well-defined topologically associating domains (TADs) (*16, 17*), chromatin compartments(*13*), and territories (*18*). For short-distance DNA interactions, we found that the contact frequencies in the trunk and viscera are much higher than those in the head. However, a reversed trend was observed for long-distance interactions, with contact frequencies in the head higher than in the trunk and viscera (Fig 2C). Additionally, the chromatin of the embryonic liver exhibited a unique conformation, with an ultra-low level of long-distance interactions but the most median-distance interactions (Kolmogorov-Smirnov test *P* = 1.18×10^-9^), as confirmed by heat maps of chromatin interactions (Fig 2D). Overall, our findings suggest a new type of distinction in hierarchical chromatin conformations (Fig 2E). Next, we explored the links to well-defined TADs, chromatin compartments, and territories, assessing their underlying mechanisms in orchestrating embryo genesis and development.

## The short-distance interactions affect embryo development

The TADs are widely reported machineries with short-distance interactions (generally at 40 kb to 3 Mb) (*17, 19*). To test if TADs vary in embryonic organs and orchestrate development-associated gene expressions, we collected the accessible data for single-cell spatial transcriptomes (*7*). In these results, 89 genes such as the core gene for brain development *Sox2*, were identified with organ-specific expressions, and exhibited frequent interactions within their 500 kb regions (Fig 3A; Table S3). Further analysis at a resolution of 40kb revealed that the bin containing *Sox2* shows a higher frequency of interactions in the brain than in other organs, like liver (Fig 3B). Given a dozen enhancer elements around *Sox2* (*20*), the frequent DNA interactions nearby may regulate embryonic brain development by strengthening the contacts of enhancers and *Sox2* (Fig 3B-C). These results explain a fundamental mechanism of how short-distance interactions affects embryo development.

**Fig. 3.**
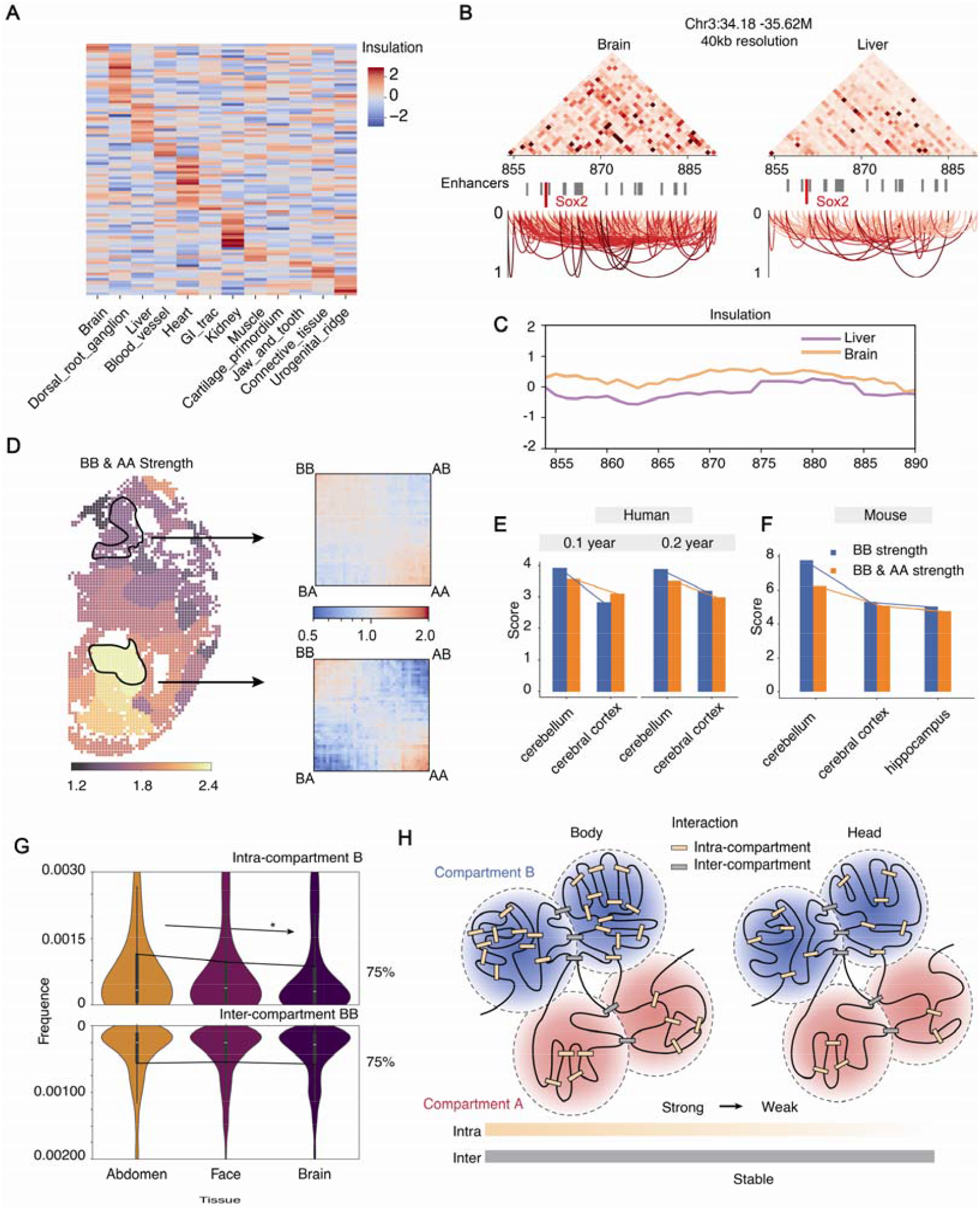
Intra-compartment interactions regulate embryo development. (**A**) Representative cross-organ marker genes insulation heatmap, where marker genes for each organ are calculated from the E12.5 mouse embryo spatial transcriptome of the MOSTA dataset. (**B**) Contact heatmap between 34.18M-35.62M of chr3 in brain and liver at 40kb resolution. (**C**) Insulation scores for spot in liver (purple) and brain (yellow). (**D**) **S**trength of compartmentalization of tissues and saddle diagram of liver and part of the brain. Compartment strength for different human (**E**) and mouse (**F**) brain areas from Dip-C. (**G**) The distribution of interaction frequency within (intra) and between (inter) compartments of multiple tissues. The color of the distribution is related to the strength of compartmentalization. (**H**) The model for interactions between and within compartments.

## Intra-but not inter-compartment interactions vary in embryonic organs

We estimated that median-distance interactions share a similar frequency in most embryonic organs (see above), which may reflect a relative stability of interactions among compartments, given that 75% of our compartments A/B exhibit less than 12Mb long. Furthermore, we calculated the strength of compartment interactions for each organ region and sub-region (Fig 3D). The interactions between DNA ends in compartments B (BB) and those in compartments A (AA) exhibit a high level of strength (Fig 3E), while those between compartments B and A (AB and BA in Fig 3E) represent a low level. This is consistent with previously reported chromatin conformation with compatibility of the same compartment type but separation between compartments A and B (*7*). However, we unexpectedly uncovered that the strengths differ in whole embryo organs and sub-organs, such as brain sub-regions; the strengths are higher in abdomen but lower in the head area (Fig 3D). This is inconsistent with the similar frequency of median-distance interactions. Meanwhile, we collected more evidence that interaction strengths of the same compartment type differ in post-birth brain sub-regions (Fig 3FG), by calculating the published data for single cell 3D genomics (*15, 21*). These results support the strength difference we uncovered, and suggest that it may initiate prior to E12.5 stage.

We proposed a conformation model to explain the conflict between the similar median-distance interactions and the distinct compartment interactions. We further divided these compartment interactions into two classes: the intra- and inter-compartment interactions. In these results, remarkable difference among organs was detected for intra-but not inter-compartment interactions (Fig 3H; Fig S7). The organic distinction in intra-compartment interactions may be consistent with the distinction for short-distance interactions, while the similarity in inter-compartment interactions coincide to the similar frequency of median-distance interactions. Taken together, our spatial 3D genomics data provides a new model suggesting that the interactions between same-type compartments share a similar frequency among embryonic organs, and only the interaction of DNA within one compartment varies and may orchestrate gene expressions for embryo development (Fig 3I).

## Embryonic liver holds an overwhelmingly packed chromatin

The long-distance interactions may affect the chromosomal territory and intra-nuclear distribution (*22*), but evidence to support their linkage to gene expressions is lacking. We uncovered that the frequency of long-distance interactions differs among embryonic organs, with the liver exhibiting an ultra-low level of this frequency (Fig 2 C and E; see above). This provides an ideal model to explore the links of chromatin conformation and gene expression. Further analysis on interactions of intra-chromosomal regions was preformed (Fig 4A), also supporting that liver has frequent median-distance interactions but fewer long-distance ones, as seen in chromosome 2. Given that cells in the embryonic liver and brain stay in identical S and G2M phases (Fig 4B), these interactions were independently of cell cycling processing. Meanwhile, 3D models of chromatin in each organ were constructed, revealing that the putative telomeric positions are at a longer distance (Fig 4C) in the liver than in the other organs, like face. This was validated by bootstrapping analysis (Fig S9). Combining these results, a unique conformation of the liver chromatin was proposed, with a low mechanical bendability limiting long-distance interactions, packed by frequent interactions between adjacent same-type compartments (Fig 4D). Additionally, we did not detect specific expression pattern in the liver. Given that liver cells have widely been reported as polyploidy or aneuploidy with a high capability of proliferation after birth (*23, 24*), our results suggest that the unique chromatin conformation may link to its special physiology, and initiates at embryo stage.

**Fig. 4.**
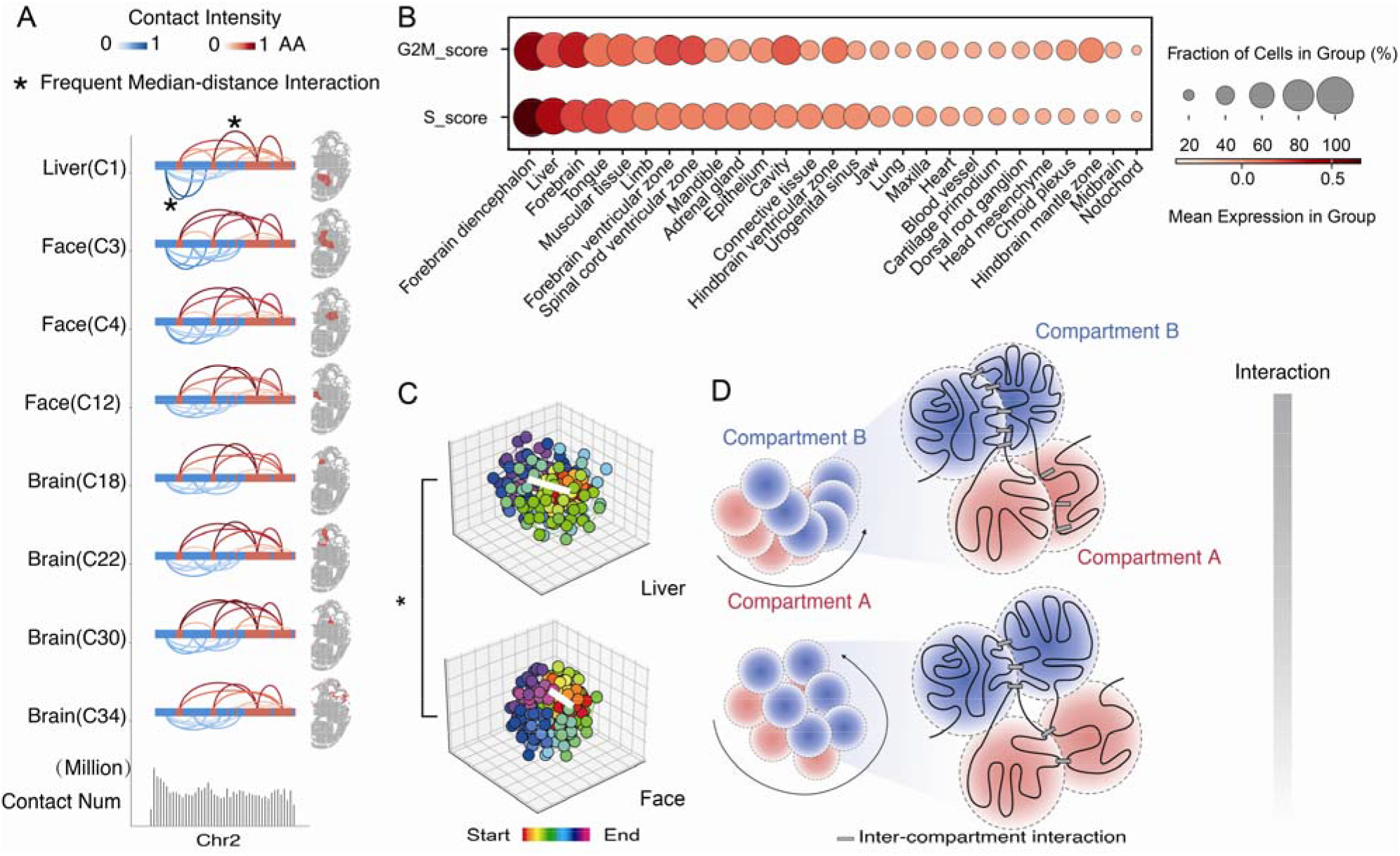
A unique conformation in the embryonic liver. (**A**) Differences in the interaction between compartments of the same type in chromosome 2. (**B**) Cell cycle scoring of different organs for E12.5 embryo. (**C**) 3D structure of chromosome 6 of the liver and face at 1 Mb resolution. (**D**) The model for a unique chromatin conformation in the liver, with enhanced adjacent-compartment interactions but few remote-compartment interactions.

## Discussion

We introduce the spatial-SPRITE approach, which assesses the chromatin conformation of nuclei *in-situ* in tissue sections with high resolution. By developing the Spatool pipeline, we constructed an atlas of hierarchical conformations for each embryonic organ. While single-cell 3D genomics technologies have been applied to the cell lines and separated cells of tissue, their data may not fully characterize the genome architecture of whole tissue due to the lack of positional information of these cells. Previous studies using single-cell 3D genomics approaches have isolated the cerebellum and cerebral cortex of human and mouse (*15*), revealing varying chromatin conformations of brain sub-regions. However, our spatial 3D genomics data detected nine sub-regions beyond the concepts of embryonic fore-, mid-, and hind-brains, suggesting that analyzing a whole tissue may yield more unbiased result results than analyzing tissue partitioned by prior knowledge. In summary, spatial 3D genomics can provide an unbiased conclusion for exploring the heterogeneity in whole tissues.

The multiway interactions from the SPRITE strategy enhance the confidence of our approach in detecting chromatin region interactions. We split the compartment interactions into the intra- and inter-compartment ones, and advanced a model suggesting that only intra-compartment interactions vary among organs to orchestrate gene expressions. Nevertheless, we also unveiled that the enhanced adjacent-compartment interactions in the liver may overwhelmingly pack chromatin with low mechanical bendability, limiting long-distance interactions. This may be linked to the liver’s special pattern for gene expression. Given that polyploid cells are common in adult liver of diverse species, potentially effecting tissue regeneration, terminal differentiation, and even tumorigenesis(*23, 24*), our data suggest that unique chromatin conformations may initiate at embryo stage. These advancements in theory and model demonstrate that our spatial 3D genomics approach is a critical tool for exploring the underlying mechanism of tissue development, particularly in the heterogeneous organs.

## Funding

This work was supported by National Key R&D Program of China (2023ZD04073) and National Natural Science Foundation of China (32350002, 32370257). X.L. was supported by the startup funding from State Key Laboratory of Plant Genomics, IGDB, CAS.

## Author contributions

X.L. supervised this work. Y.M. designed and completed the experimental procedure of Spatial-SPRITE. B.G. and Y.X. developed a pipeline to process data. M.S. and F.L. provided the mouse material. X.L., Y.M., B.G., and Y.X. prepared the manuscript.

## Competing interests

The Spatial-SPRITE procedure and its applications are covered in a pending patent.

## Data and materials availability

The raw sequence data reported in this paper have been deposited in the Genome Sequence Archive (Genomics, Proteomics & Bioinformatics 2021) in National Genomics Data Center (Nucleic Acids Res 2022), China National Center for Bioinformation / Beijing Institute of Genomics, Chinese Academy of Sciences (GSA: CRA014655) that are publicly accessible at https://ngdc.cncb.ac.cn/gsa.

## References

1. B. Bonev, G. Cavalli, Organization and function of the 3D genome. Nature Reviews Genetics 17, 661–678 (2016).

2. T. Zhou, R. Zhang, J. Ma, The 3D Genome Structure of Single Cells. Annu Rev Biomed Data Sci 4, 21–41 (2021).

3. I. M. Flyamer, J. Gassler, M. Imakaev, H. B. Brandao, S. V. Ulianov, N. Abdennur, S. V. Razin, L. A. Mirny, K. Tachibana-Konwalski, Single-nucleus Hi-C reveals unique chromatin reorganization at oocyte-to-zygote transition. Nature 544, 110-+ (2017).

4. T. Nagano, Y. Lubling, C. Varnai, C. Dudley, W. Leung, Y. Baran, N. M. Cohen, S. Wingett, P. Fraser, A. Tanay, Cell-cycle dynamics of chromosomal organization at single-cell resolution. Nature 547, 61-+ (2017).

5. (!!! INVALID CITATION !!! (5-11)).

6. S. G. Rodriques, R. R. Stickels, A. Goeva, C. A. Martin, E. Murray, C. R. Vanderburg, J. Welch, L. M. Chen, F. Chen, E. Z. Macosko, Slide-seq: A scalable technology for measuring genome-wide expression at high spatial resolution. Science 363, 1463-+ (2019).

7. A. Chen, S. Liao, M. Cheng, K. Ma, L. Wu, Y. Lai, X. Qiu, J. Yang, J. Xu, S. Hao, X. Wang, H. Lu, X. Chen, X. Liu, X. Huang, Z. Li, Y. Hong, Y. Jiang, J. Peng, S. Liu, M. Shen, C. Liu, Q. Li, Y. Yuan, X. Wei, H. Zheng, W. Feng, Z. Wang, Y. Liu, Z. Wang, Y. Yang, H. Xiang, L. Han, B. Qin, P. Guo, G. Lai, P. Muñoz-Cánoves, P. H. Maxwell, J. P. Thiery, Q.-F. Wu, F. Zhao, B. Chen, M. Li, X. Dai, S. Wang, H. Kuang, J. Hui, L. Wang, J.-F. Fei, O. Wang, X. Wei, H. Lu, B. Wang, S. Liu, Y. Gu, M. Ni, W. Zhang, F. Mu, Y. Yin, H. Yang, M. Lisby, R. J. Cornall, J. Mulder, M. Uhlén, M. A. Esteban, Y. Li, L. Liu, X. Xu, J. Wang, Spatiotemporal transcriptomic atlas of mouse organogenesis using DNA nanoball-patterned arrays. Cell, (2022).

8. Y. Liu, M. Yang, Y. Deng, G. Su, A. Enninful, C. C. Guo, T. Tebaldi, D. Zhang, D. Kim, Z. Bai, E. Norris, A. Pan, J. Li, Y. Xiao, S. Halene, R. Fan, High-Spatial-Resolution Multi-Omics Sequencing via Deterministic Barcoding in Tissue. Cell 183, 1665–1681 e1618 (2020).

9. Y. Deng, M. Bartosovic, P. Kukanja, D. Zhang, Y. Liu, G. Su, A. Enninful, Z. Bai, G. Castelo-Branco, R. Fan, Spatial-CUT&Tag: Spatially resolved chromatin modification profiling at the cellular level. Science 375, 681–686 (2022).

10. Y. Deng, M. Bartosovic, S. Ma, D. Zhang, P. Kukanja, Y. Xiao, G. Su, Y. Liu, X. Qin, G. B. Rosoklija, A. J. Dwork, J. J. Mann, M. L. Xu, S. Halene, J. E. Craft, K. W. Leong, M. Boldrini, G. Castelo-Branco, R. Fan, Spatial profiling of chromatin accessibility in mouse and human tissues. Nature, (2022).

11. S. A. Quinodoz, N. Ollikainen, B. Tabak, A. Palla, J. M. Schmidt, E. Detmar, M. M. Lai, A. A. Shishkin, P. Bhat, Y. Takei, V. Trinh, E. Aznauryan, P. Russell, C. Cheng, M. Jovanovic, A. Chow, L. Cai, P. McDonel, M. Garber, M. Guttman, Higher-Order Inter-chromosomal Hubs Shape 3D Genome Organization in the Nucleus. Cell 174, 744-757.e724 (2018).

12. J. Dekker, A. S. Belmont, M. Guttman, V. O. Leshyk, J. T. Lis, S. Lomvardas, L. A. Mirny, C. C. O’Shea, P. J. Park, B. Ren, J. C. R. Politz, J. Shendure, S. Zhong, A. van den Berg, A. B. Heckert, A. Bertero, A. Bortnick, A. Kukalev, A. Moore, A. Pombo, A. S. Hansen, A. M. Chiariello, A. Sali, A. Belmont, A. Stephens, A. Nand, A. L. Valton, A. Goloborodko, A. He, B. van Steensel, B. Webb, B. Roscoe, B. Li, B. Ren, B. Chait, C. A. Blau, C. Annunziatella, C. Ware, C. L. Wei, C. Leemans, C. Disteche, C. Jarjour, C. Thieme, C. Murry, C. T. Barcia, C. Trapnell, C. Murre, D. Peric-Hupkes, D. Simon, D. Bartlett, D. Gao, D. Plewczynski, D. Gilbert, D. Gorkin, D. McSwiggen, D. Lin, D. Aghamirzaie, E. Banigan, E. Finn, E. Sontheimer, F. T. Cadete, F. Alber, F. Mast, G. Filippova, G. G. Yardimci, G. Fudenberg, G. Loof, G. Bonora, G. Pegoraro, G. Caglio, G. Polles, H. Ozadam, H. Shin, H. Pliner, H. Reinecke, H. C. Li, H. Tjong, H. Fang, H. Marie-Nelly, H. Belaghzal, H. Brandao, H. M. Zhao, I. Cisse, I. Y. Jung, I. Tasan, I. Juric, J. O. Andrews, J. Schreiber, J. H. Spille, J. Zimmerman, J. Shendure, J. Dixon, J. Ma, J. Xu, J. Sima, J. Dekker, J. Gibcus, J. Nuebler, J. Aitchison, J. Marko, J. Lam, J. A. B. Mendieta, J. C. R. Mulia, J. Cayford, K. Cook, K. Mitzelfelt, K. M. Parsi, K. Klein, L. Brueckner, L. Mirny, L. Zhang, L. Pabon, L. Chen, L. Carpp, L. Y. Yang, L. Pei, M. Sander, M. Imakaev, M. Nicodemi, M. Schueler, M. Falk, M. Denholtz, M. Libbrecht, M. F. Bolukbasi, M. Zhen, M. Yu, M. Rout, M. Hu, M. Mir, N. Armani, N. Hua, N. Kubo, N. Abdennur, N. Krietenstein, N. Khanna, O. Dudko, O. Rando, O. Luo, P. Chaturvedi, P. Blainey, P. Fields, P. Wang, Q. J. Li, R. Casellas, R. Gudla, R. Maeh, R. Kempfer, R. Beagrie, R. Biggs, R. X. Fang, R. L. Qiu, R. M. J. Genga, S. Srivatsan, S. Kumar, S. Wolfe, S. Shaffer, S. S. Kim, S. Shachar, S. Bianco, S. Jain, T. Sasaki, T. Isoda, T. Misteli, T. van Schaik, T. Liu, T. H. Hsieh, V. Ramani, V. Agarwal, V. Dileep, V. Chandra, W. Winick-Ng, W. Y. Li, W. Noble, X. Darzacq, X. H. J. Zhou, X. X. Deng, X. Xiong, X. L. Yang, Y. Yang, Y. Zhang, Y. Kou, Y. Zhou, Y. J. Ruan, Y. Chen, Y. C. Wang, Y. J. Qiu, Z. J. Duan, Z. H. Tang, A. Ozer, A. Cote, A. Tanay, A. Chow, A. D. Omer, A. Hwang, C. Dudley, C. Bartman, C. Danko, C. Varnai, E. L. Aiden, G. Blobel, H. N. Lin, J. Phillips-Cremins, J. Lis, J. Wang, J. Ray, M. Dunagin, M. Arrastia, M. Lai, M. Curtis, M. Kushner, M. Pham, M. Wang, M. Yang, M. Guttman, N. C. Durand, N. Ollikainen, P. Munn, P. Fraser, R. Ismagilov, S. Hsu, S. Bhardwaj, S. Quinodoz, T. Nagano, T. Amarante, W. Zipfel, Y. Baran, Y. Lubling, Z. Wang, A. Palla, A. Muimbey-Wahula, A. Vertii, A. Moradian, C. Larabell, C. Brangwynne, C. Lindsay, D. Sanders, D. Scalzo, E. Cannavo, G. McDermott, H. Ozadam, H. Ma, J. Moresco, J. Ritland, J. Dekker, J. Rinn, J. Yates, J. Zhu, K. Roth, L. Gerace, L. Tait, L. Brown, L. Zhu, M. Kordon, M. Groudine, M. Le Gros, M. Escamilla, M. Sweredoski, M. Guttman, P. Kaufman, P. Maas, R. Barutcu, R. Amin, S. Baboo, S. M. Debartolome, S. Hess, S. Lomvardas, T. Pederson, T. Szempruch, W. Walkup, X. M. Sun, Y. D. Shin, A. Senecal, A. Hansen, A. Barentine, A. Spakowitz, A. K. Gustavsson, A. Tangara, B. Rieger, B. Nijmeijer, B. Lim, B. English, C. Barton, C. Kenworthy, C. Carroll, C. O’Shea, D. Boassa, D. Baddeley, D. Grunwald, E. Birney, F. Chuang, G. Castillon, H. F. Wang, H. Grabmayr, H. T. Chen, H. Ou, J. Ellenberg, J. Liphardt, J. Soroczynski, J. Biswas, J. Yao, J. W. Yin, J. Bewersdorf, J. Ries, J. Bardales, J. Roberti, K. Zaret, K. Chung, K. Lam, L. S. Qi, L. Schmitt, L. Barinov, L. C. Tu, L. L. Yang, L. Tian, L. Cai, M. Ellisman, M. Mackey, M. Haberl, M. Huisman, M. Clark, M. Levo, M. Levine, M. Mir, N. Walther, O. Oedegaard, P. Guo, Q. S. Zheng, R. H. Cheng, R. Ghosh, R. Ramachandra, R. Coleman, R. Singer, R. W. Liu, R. Walden, S. Phan, S. Ramachandra, R. Coleman, R. Singer, R. W. Liu, R. Walden, S. Phan, S. Quanming, S. Ganguly, S. Alexander, S. Peltier, T. Fukaya, T. Deerinck, T. Gregor, T. Fitzgerald, W. Moerner, X. Darzacq, Y. D. Zhang, Y. M. Li, Y. Takei, Y. Izumiya, Y. Lin, Z. Frankenstein, B. Ren, C. Kling, C. Rivera, H. Z. Zheng, K. Z. Rivera, L. Hebert, M. Rivas-Astroza, Q. Y. Wu, R. Calandrelli, S. Subramaniam, S. Zhong, S. Chien, V. Leshyk, W. Z. Chen, X. Y. Cao, Z. M. Yan, A. Balashov, A. Schroeder, A. Goloborodko, B. H. Alver, C. Vitzthum, C. Nam, D. F. Li, D. Purushotham, E. C. Pehrsson, F. Yue, F. Lekschas, H. Pfister, H. Strobelt, H. Brandao, H. S. Jang, J. Luber, J. Hwang, J. Walsh, J. Johnson, J. Nubler, K. Kirli, L. Mirny, M. Falk, M. Imakaev, M. N. Choudhary, N. Abdennur, N. Gehlenborg, P. Kerpedjiev, P. Park, P. V. Kharchenko, R. L. Sears, S. Lee, S. Wang, T. Yang, T. X. M. Hu, T. Wang, Y. Hou, D. N. Network, The 4D nucleome project. Nature 549, 219–226 (2017).

13. E. Lieberman-Aiden, N. L. van Berkum, L. Williams, M. Imakaev, T. Ragoczy, A. Telling, I. Amit, B. R. Lajoie, P. J. Sabo, M. O. Dorschner, R. Sandstrom, B. Bernstein, M. A. Bender, M. Groudine, A. Gnirke, J. Stamatoyannopoulos, L. A. Mirny, E. S. Lander, J. Dekker, Comprehensive mapping of long-range interactions reveals folding principles of the human genome. Science 326, 289–293 (2009).

14. J. P. Fortin, K. D. Hansen, Reconstructing A/B compartments as revealed by Hi-C using long-range correlations in epigenetic data. Genome Biol 16, 180 (2015).

15. L. Tan, J. Shi, S. Moghadami, B. Parasar, C. P. Wright, Y. Seo, K. Vallejo, I. Cobos, L. Duncan, R. Chen, K. Deisseroth, Lifelong restructuring of 3D genome architecture in cerebellar granule cells. Science 381, 1112–1119 (2023).

16. T. Sexton, E. Yaffe, E. Kenigsberg, F. Bantignies, B. Leblanc, M. Hoichman, H. Parrinello, A. Tanay, G. Cavalli, Three-dimensional folding and functional organization principles of the Drosophila genome. Cell 148, 458–472 (2012).

17. J. R. Dixon, S. Selvaraj, F. Yue, A. Kim, Y. Li, Y. Shen, M. Hu, J. S. Liu, B. Ren, Topological domains in mammalian genomes identified by analysis of chromatin interactions. Nature, (2012).

18. J. A. Croft, J. M. Bridger, S. Boyle, P. Perry, P. Teague, W. A. Bickmore, Differences in the localization and morphology of chromosomes in the human nucleus. J Cell Biol 145, 1119–1131 (1999).

19. R. Kempfer, A. Pombo, Methods for mapping 3D chromosome architecture. Nature Reviews Genetics 21, 207–226 (2020).

20. D. U. Gorkin, I. Barozzi, Y. Zhao, Y. Zhang, H. Huang, A. Y. Lee, B. Li, J. Chiou, A. Wildberg, B. Ding, B. Zhang, M. Wang, J. S. Strattan, J. M. Davidson, Y. Qiu, V. Afzal, J. A. Akiyama, I. Plajzer-Frick, C. S. Novak, M. Kato, T. H. Garvin, Q. T. Pham, A. N. Harrington, B. J. Mannion, E. A. Lee, Y. Fukuda-Yuzawa, Y. He, S. Preissl, S. Chee, J. Y. Han, B. A. Williams, D. Trout, H. Amrhein, H. Yang, J. M. Cherry, W. Wang, K. Gaulton, J. R. Ecker, Y. Shen, D. E. Dickel, A. Visel, L. A. Pennacchio, B. Ren, An atlas of dynamic chromatin landscapes in mouse fetal development. Nature, (2020).

21. L. Tan, W. Ma, H. Wu, Y. Zheng, D. Xing, R. Chen, X. Li, N. Daley, K. Deisseroth, X. S. Xie, Changes in genome architecture and transcriptional dynamics progress independently of sensory experience during post-natal brain development. Cell 184, 741-+ (2021).

22. T. Nagano, Y. Lubling, T. J. Stevens, S. Schoenfelder, E. Yaffe, W. Dean, E. D. Laue, A. Tanay, P. Fraser, Single-cell Hi-C reveals cell-to-cell variability in chromosome structure. Nature 502, 59-+ (2013).

23. J. E. Guidotti, O. Bregerie, A. Robert, P. Debey, C. Brechot, C. Desdouets, Liver cell polyploidization: A pivotal role for binuclear hepatocytes. Journal of Hepatology, (2003).

24. R. Donne, M. Saroul-Aïnama, P. Cordier, S. Celton-Morizur, C. Desdouets, Polyploidy in liver development, homeostasis and disease. Nature Reviews Gastroenterology & Hepatology, (2020).

